# Gene editing of the E3 ligase *PIRE1* fine-tunes ROS production for enhanced bacterial disease resistance in tomato

**DOI:** 10.1101/2024.07.31.606097

**Authors:** Bardo Castro, Suji Baik, Megann Tran, Jie Zhu, Tianrun Li, Andrea Tang, Nathalie Aoun, Alison C Blundell, Michael Gomez, Elaine Zhang, Myeong-Je Cho, Tiffany Lowe-Power, Shahid Siddique, Brian Staskawicz, Gitta Coaker

## Abstract

Reactive oxygen species (ROS) accumulation is required for effective plant defense. Accumulation of the Arabidopsis NADPH oxidase RBOHD is regulated by phosphorylation of a conserved C-terminal residue (T912) leading to ubiquitination by the RING E3 ligase PIRE. Arabidopsis *PIRE* knockouts exhibit enhanced ROS production and resistance to the foliar pathogen *Pseudomonas syringae*. Here, we identified 170 *PIRE* homologs, which emerged in Tracheophytes and expanded in Angiosperms. We investigated the role of *Solanum lycopersicum* (tomato) PIRE homologs in regulating ROS production, RBOH stability, and disease resistance. Mutational analyses of residues corresponding to T912 in the tomato RBOHD ortholog, SlRBOHB, affected protein accumulation and ROS production in a *PIRE-*dependent manner. Using CRISPR-cas9, we generated mutants in two *S. lycopersicum PIRE* homologs (*SlPIRE*). *SlPIRE1* edited lines (*Slpire1*) in the tomato cultivar M82 displayed enhanced ROS production upon treatment with flg22, an immunogenic epitope of flagellin. Furthermore*, Slpire1* exhibited decreased disease symptoms and bacterial accumulation when inoculated with foliar bacterial pathogens *Pseudomonas syringae* and *Xanthomonas campestris*. However, *Slpire1* exhibited similar levels of colonization as wild type upon inoculation with diverse soilborne pathogens. These results indicate that phosphorylation and ubiquitination crosstalk regulate RBOHs in multiple plant species, and *PIRE* is a promising target for foliar disease control. This study also highlights the pathogen-specific role of *PIRE*, indicating its potential for targeted manipulation to enhance foliar disease resistance without affecting root-associated interactions, positioning *PIRE* as a promising target for improving overall plant health.

## Introduction

Crop production is impacted by diverse plant pathogens. Among five major food crops (potato, soybean, wheat, maize, and rice) losses due to pests and pathogens range between 17% and 30% globally (Savary et al., 2019). Plants contain innate immune receptors that can recognize all pathogen classes. Pathogen recognition can occur extracellularly via cell-surface localized pattern recognition receptors (PRRs) leading to pattern-triggered immunity (PTI), or intracellularly through recognition of pathogen encoded effectors by nucleotide-binding domain leucine-rich repeat receptors (NLRs) leading to effector-triggered immunity (ETI) (Yuan et al., 2023). NLRs and PRRs mutually potentiate each other and their activation leads to convergent responses (Yuan et al., 2023). Common plant immune responses include ion influxes, rapid production of reactive oxygen species (ROS), transcriptional reprogramming, deposition of structural barriers, and stomatal closure, all of which culminate in resistance (Yuan et al., 2023).

Much of our understanding of PTI comes from the conserved PRR, FLAGELLIN-SENSING 2 (FLS2). FLS2 is a leucine-rich repeat receptor kinase (LRR-RK) perceives a 22 amino acid immunogenic epitope, flg22, from the bacterial flagellin protein FliC (Zipfel et al., 2004). The molecular interaction between flg22 and FLS2 leads to recruitment of a SERK (somatic embryogenesis receptor kinase) co-receptor (Chinchilla et al., 2007; Heese et al., 2007; Sun et al., 2013). In Arabidopsis, formation of the FLS2 receptor complex induces trans-phosphorylation of multiple intracellular kinases, including receptor-like cytoplasmic kinases, calcium-dependent protein kinases, and mitogen-activated protein kinases (MAPKs), which lead to multiple defense outputs (Couto & Zipfel, 2016). To rapidly respond to pathogens, plant immune receptors and key signaling proteins are pre-synthesized and regulated through post-translational modifications (PTMs). PTMs can affect all aspects of protein function including dynamic control or protein abundance, activity, and localization (Csizmok & Forman-Kay, 2018; Lee et al., 2023). For instance, the FLS2 receptor complex, as well as calcium and ROS production are regulated through multiple transphosphorylation events (Couto & Zipfel, 2016; Kadota et al., 2014; Li et al., 2014; Thor et al., 2020; Tian et al., 2019; Zhang et al., 2018). Another key layer of post-translational regulation is ubiquitination and subsequent degradation. For example, after the FLS2-flg22 immune complex forms, it is ubiquitinated by two U-box E3 ubiquitin ligases, PUB12 and PUB13, leading to its degradation and immune signal turnover (Lu et al., 2011).

One pivotal process regulated by phosphorylation is the production of apoplastic ROS by membrane localized NADPH oxidases, termed respiratory burst oxidase homologs (RBOHs) in plants (Castro et al., 2021). RBOHs produce superoxide (O^2•−^), which can be converted to hydrogen peroxide (H_2_O_2_), which is the most stable form and considered a key signaling molecule (Castro et al., 2021). RBOH activation during PTI leads to rapid and dynamic generation of ROS. Extracellular accumulation of ROS are involved in numerous processes including cell wall lignification, stomatal closure, and systemic acquired resistance (Kadota et al., 2015; Waszczak et al., 2018). Although de novo ROS production is crucial for defense, continual accumulation of hydrogen peroxide, superoxide, and hydroxyl radicals can lead to cellular oxidative damage (Kerchev & Van Breusegem, 2022).

The production of ROS is essential for a robust immune response; however, this production must be dynamically regulated to minimize detrimental effects to the host. During pathogen perception, different kinase families phosphorylate N-terminal residues on Arabidopsis RBOHD (AtRBOHD), leading to functional activation (Bender & Zipfel, 2023; Kadota et al., 2015; Zhang et al., 2018). In recent years, research has shown that modification of C-terminal residues of AtRBOHD are also important for its regulation (Kimura et al., 2020; Lee et al., 2020). Our previous work identified the receptor-like cytoplasmic kinase PBS1-like kinase 13 (PBL13) that phosphorylates multiple AtRBOHD C-terminal residues to negatively regulate ROS production (Lee et al., 2020). PBL13 phosphorylates T912, which reduces AtRBOHD stability, and S862, which impacts enzyme activity (Lee et al., 2020). Crosstalk between phosphorylation and ubiquitination is critical to dynamically control protein levels (Castro et al., 2021; Lu et al., 2011; Swaney et al., 2013). The PBL13 interacting RING domain E3 ligase (PIRE) ubiquitinates AtRBOHD’s C-terminus in a phosphorylation dependent manner (D. Lee et al., 2020). Consistent with these results, *pbl13* and *pire* knockouts displayed enhanced AtRBOHD accumulation, immune-induced ROS production, and resistance to the bacterial pathogen *Pseudomonas syringae* (Lee et al., 2020). Upon pathogen perception, phosphatidic acid binds to RBOHD, inhibiting its interaction with PIRE (Qi et al., 2024).This suppression prevents RBOHD protein degradation, resulting in increased levels of RBOHD in the plasma membrane during pathogen perception (Qi et al., 2024).

Analysis of 112 plant RBOH homologs revealed high conservation of residue T912, which is important for PBL13-PIRE regulation (Castro et al., 2021). However, PBL13 is only found in the *Brassicacea* (Lee et al., 2020). The conservation of RBOHD T912 indicates other plants may regulate RBOHs in a similar manner, but through different kinases, which can be exploited for disease control. In this manuscript we have identified homologs of the Arabidopsis ubiquitin E3 ligase PIRE across the plant kingdom and investigated the importance of RBOH modification in the Solanaceae. We investigated the importance of *Solanum lycopersicum* (tomato) and Nicotiana *PIRE* homologs for regulation of ROS activity, utilizing the *S. lycopersicum* ortholog of AtRBOHD, SlRBOHB (Li et al., 2015). SlRBOHB abundance is also regulated at similar residues and silencing in *N. benthamiana* implicated *PIRE* homologs in regulating RBOH abundance. Furthermore, we utilized CRISPR-Cas9 to generate *S. lycopersicum pire* mutants. The *S. lycopersicum pire1* mutant exhibited higher ROS production upon immune activation and increased disease resistance to foliar bacterial pathogens. Our results provide evidence that crosstalk between phosphorylation and ubiquitination functions as a conserved regulatory module for plant RBOHs and *PIRE* is a promising target to enhance disease resistance.

## Results

### Homologs of the E3 ubiquitin ligase PIRE are broadly conserved

Analysis of 112 RBOH homologs in plants revealed that the residue corresponding to T912 is highly conserved, indicating that PIRE-mediated regulation of ROS may also be conserved (Castro et al., 2021). Therefore, we first sought to identify PIRE homologs in various plant lineages. Previously, Arabidopsis RING domain proteins were classified into eight different classes based on their metal ligand residues (Stone et al., 2005). While there are more than 470 RING E3 ligases in Arabidopsis, there are only 10 identified zinc-binding RING-C2’s (Cho et al., 2017; Deshaies & Joazeiro, 2009; Duplan & Rivas, 2014; Metzger et al., 2014). AtPIRE is 319 amino acids (aa) in length and contains a modified RING-C2 domain on its C-terminal region from aa 244 to 290. The modified RING-C2 domain in Arabidopsis contains variable regions between the specific ligand binding sites (**Supplementary Fig. S1**). Utilizing SMART (Simple Modular Architecture Research Tool) (Letunic et al., 2021), we identified a low complexity region containing serine (S) and glutamic acid (D) repeats from aa 117 to 159 in AtPIRE.

Next, we investigated the emergence of AtPIRE homologs across different algal and plant lineages. We required AtPIRE homologs to possess both the low complexity region and the C-terminally localized modified RING-C2 domain (**Supplementary Fig. S1**). Using a combination of BLASTP based on the RING-C2 domain coupled with the presence of the low complexity region, we identified 170 different PIRE homologs across 64 plant species (**Supplementary Dataset 1)**. These homologs contain highly conserved in the modified RING-C2 domain region, which is important for zinc binding (**Supplementary Fig. S1, Fig. 1B**). RING proteins identified *D. salina,* in the phylum Chlorophyta (Green Algae) have N-terminal localized RING domains, but this domain does not contain the modified RING-C2 zinc-binding regions (**Supplementary Fig. S1, Fig. 1A**). Interestingly, *Chara braunii*, a member of the phylum Charophyta that emerged later than Chlorophyta, displayed a C-terminal localized ring domain, however this domain does not have all the modified RING-C2 residues (**Supplementary Fig. S1, Fig. 1A**). PIRE-like architecture was also identified in Bryophyta, but members also lacked full RING-C2 residues (**Supplementary Fig. S1**). It was not until gymnosperms that both the PIRE architecture and complete modified RING-C2 residues appeared (**Supplementary Fig. S1**). PIRE homologs significantly expanded in angiosperms, and we were able to identify members in all analyzed monocot and dicot genomes (**Fig. 1, Supplementary Table S1)**. Our analysis revealed that the complete PIRE protein architecture likely arose in gymnosperms.

**Figure 1.**
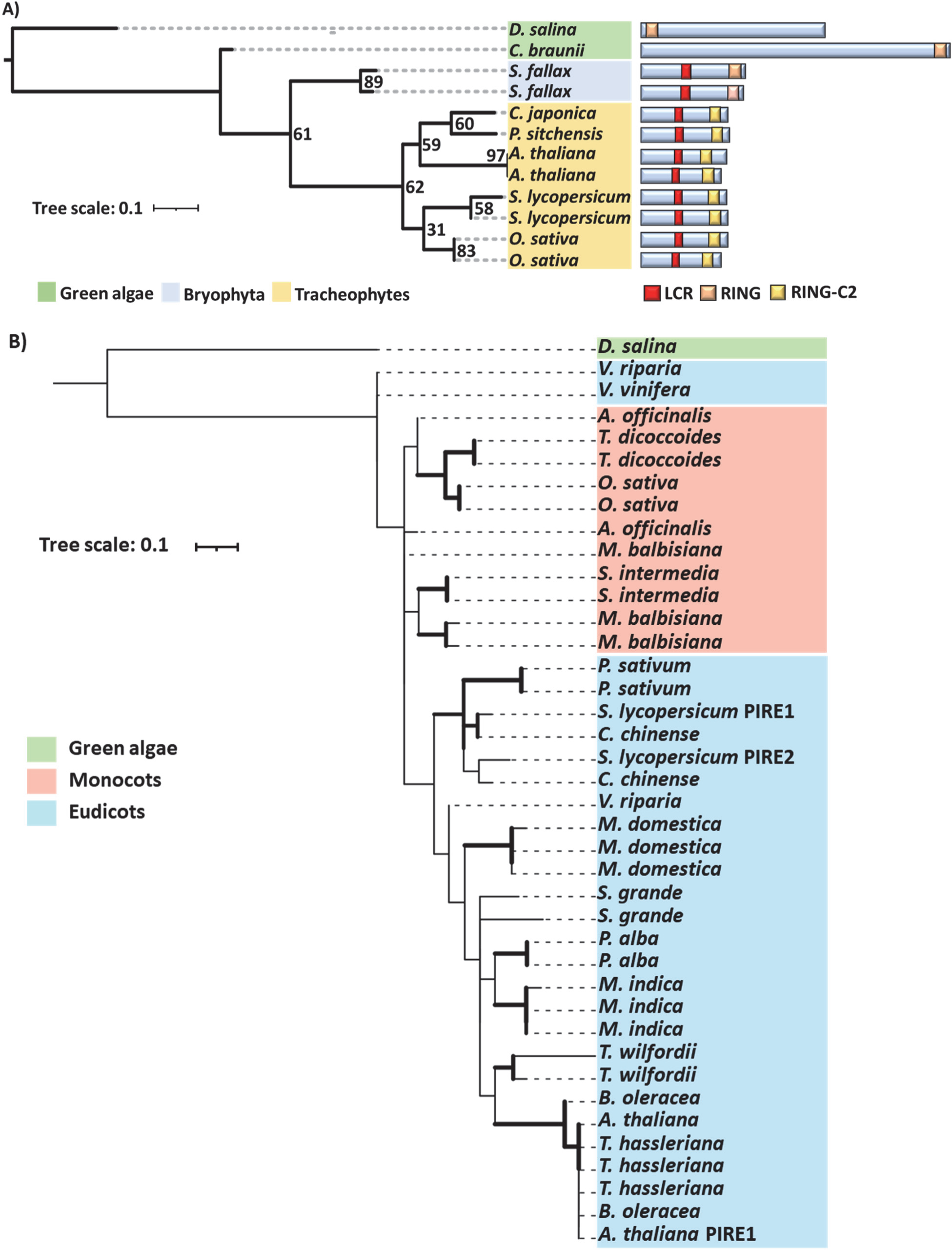
Homologs of the RING E3 ligase PIRE are present in the Tracheophytes and expanded in angiosperms. **A)** PIRE homologs are detected in the Tracheophytes. Phylogeny of the RING domain from *Arabidopsis* PIRE and closest homologs throughout the plant kingdom. The phylogenetic tree was generated using the maximum likelihood method with a bootstrap value of 1000 using IQtree. Right: Domain architecture of PIRE homologs, which contain a C-terminal modified RING-C2 domain, and a low complexity region (LCR) enriched in serine and glutamic acid residues in the central region of the protein. **B)** Phylogeny of the RING domain from 39 PIRE protein homologs identified in 20 different plant species. The phylogenetic tree was generated using the maximum likelihood method with a bootstrap value of 1000. Sequences alignments were generated utilizing Clustal Omega. Branches supported with bootstrap values above 70 have increased thickness.

### A conserved C-terminal RBOH residue regulates ROS production and abundance

Given the conservation of PIRE as well as the phosphorylated C-terminal residue corresponding to T912 in AtRBOHD (Lee et al., 2020), we sought to determine if additional pant NADPH oxidases are similarly impacted by the presence of this conserved residue. To this end, we investigated the *S. lycopersicum* SlRBOHB, which is the closest *S. lycopersicum* homolog to AtRBOHD and has previously been linked with ROS production upon flg22 perception (Li et al., 2015). AtRBOHD residue T912 corresponds to SlRBOHB T856 (**Fig. 2A**). To validate the importance of T856 in regulating ROS production and stability of SlROHB, we generated both a phosphonull (SlRBOHB^T856A^) and phosphomimic (SlRBOHB^T856D^) mutants of SlRBOHB fused to an epitope tag (YFP). We then transiently expressed these phosphomutants in *Nicotiana benthamiana* and induced PTI through flg22 treatment to measure production of ROS. Flg22 induced ROS was detected during the empty vector (EV) treatment due to endogenous RBOHB in *N. benthamiana* (**Supplementary Fig. S2**). ROS produced upon treatment with flg22 after transient expression of SlRBOHB was 10-fold higher than the EV control, demonstrating we can use this assay to detect alterations in ROS after expression of additional RBOHs (**Supplementary Fig. S2**).

**Figure 2.**
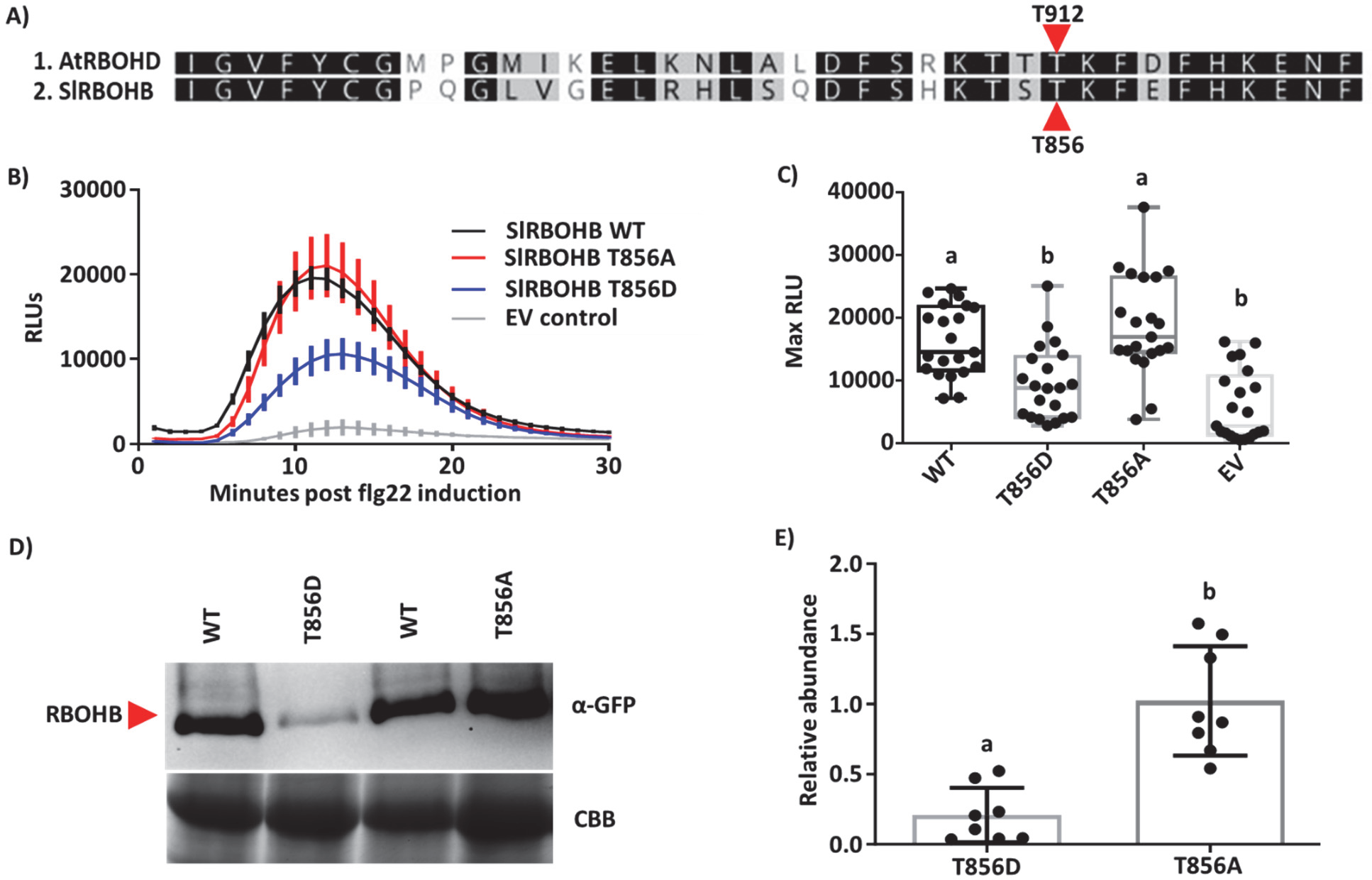
Mutations in conserved C-terminal residues *Solanum lycopersicum* RBOHB lead to changes in ROS production and protein accumulation. **A)** C-terminal amino acid alignment of the NADPH oxidases from Arabidopsis (AtRBOHD) and *S. lycopersicum* (SlRBOHB). The previously identified phosphorylated threonine 912 (T912) in AtRBOHD corresponds to threonine 856 (T856) in SlRBOHB. **B)** Different RBOHB variants were transiently expressed in *Nicotiana benthamiana.* Leaf disks were collected from *N. benthamiana* and treated with 100nM flg22 to induce ROS production over 30 minutes. Results display the mean ±SE, n=7 leaf disks. Phosphomimetic SlRBOHB^T856D^ has decreased production of reactive oxygen species (ROS) compared to SlRBOHB^WT^ and SlRBOHB^T856A^. **C)** SlRBOHB^T856D^ ROS production is significantly lower than SlRBOHB^WT^ and SlRBOHB^T856A^ post-flg22 induction as described above. Results display maximum relative light units (max RLU) of 3 independent experiments (n=21 plants). Whiskers show minimum and maximum values. Statistical differences were determined by ANOVA with post-hoc Tukey test (p value < 0.0001). **D)** SlRBOHB protein abundance was visualized by anti-GFP immunoblot 48h post-transient expression in *N. benthamiana*. SlRBOHB^T856D^ displayed reduced accumulation in *N. benthamiana* compared to SlRBOHB^WT^ and SlRBOHB^T856A^. **E)** SlRBOHB protein accumulation was quantified from anti-GFP immunoblots utilizing Image Lab. Protein levels were first normalized using the rubisco band from the Coomassie brilliant blue (CBB) gel, then the relative intensity of each protein was compared to SlRBOHB^WT^. N=8, error bars represent standard deviation (SD). Statistical differences were calculated by a Kruskal-Wallis test with a Dunn’s test (p value = 0.0003). SlRBOHB^T856D^ has significantly lower protein accumulation than SlRBOHB^WT^ and SlRBOHB^T856A^.

Transient expression of wild-type SlRBOHB (SlRBOHB^WT^) and SlRBOHB^T856A^ led to similar levels of ROS production post flg22 induction. However, transient expression of SlRBOHB^T856D^ led to a significant decrease in ROS production post flg22 induction (**Fig. 2B-C**). Although there was a decrease in ROS production for SlRBOHB^T856D^, we did not detect changes in the temporal dynamics of ROS production with all transiently expressed SlRBOHB variants (**Fig. 2B**).

In Arabidopsis, phosphorylation of T912 leads to vacuolar degradation of AtRBOHD (D. Lee et al., 2020). Therefore, we hypothesized that decreased ROS production was due to reduced accumulation of the phosphomimic SlRBOHB^T856D^. We quantified the accumulation SlRBOHB and respective phosphomutants by western blot after transient expression in *N. benthamiana*. The accumulation of YFP-tagged SlRBOHB^WT^ and SlRBOHB^T856A^ were not significantly different from one another (**Fig. 2D, E**). However, SlRBOHB^T856D^ displayed decreased accumulation by immunoblot analyses when compared to both SlRBOHB^WT^ and SlRBOHB^T856A^ (**Fig. 2D, E**). These results are consistent with the regulation of AtRBOHD in Arabidopsis, where phosphorylation of T912 led to enhanced degradation of AtRBOHD (Lee et al., 2020). In our case, phosphomimic mutations of the corresponding residue in SlRBOHB, T856, also led to decreased accumulation of SlRBOHB during transient expression and in turn reduced production of ROS. These findings further support that phosphorylation of conserved residues play an essential role in regulating NADPH abundance and ROS production.

### Accumulation of the SlRBOHB T856 phosphomimic is dependent on *PIRE* homologs

Next, we sought to determine if the abundance of SlRBOHB is dependent on *PIRE* homologs. There are two *S. lycopersicum PIRE* homologs, *SlPIRE1* and *SlPIRE2* (**Fig. 1A**). Using amino acid sequence alignments, we generated a phylogenetic tree to identify *N. benthamiana* homologs of *SlPIRE1* and *SlPIRE2*. Utilizing this method we identified five homologs in *N. benthamiana*: three homologs of *SlPIRE1* (*NbPIRE 1-1, NbPIRE 1-2, and NbPIRE 1-3*) and two homologs of *SlPIRE2* (*NbPIRE 2-1 and NbPIRE 2-2*) (**Fig. 3A**).

**Figure 3.**
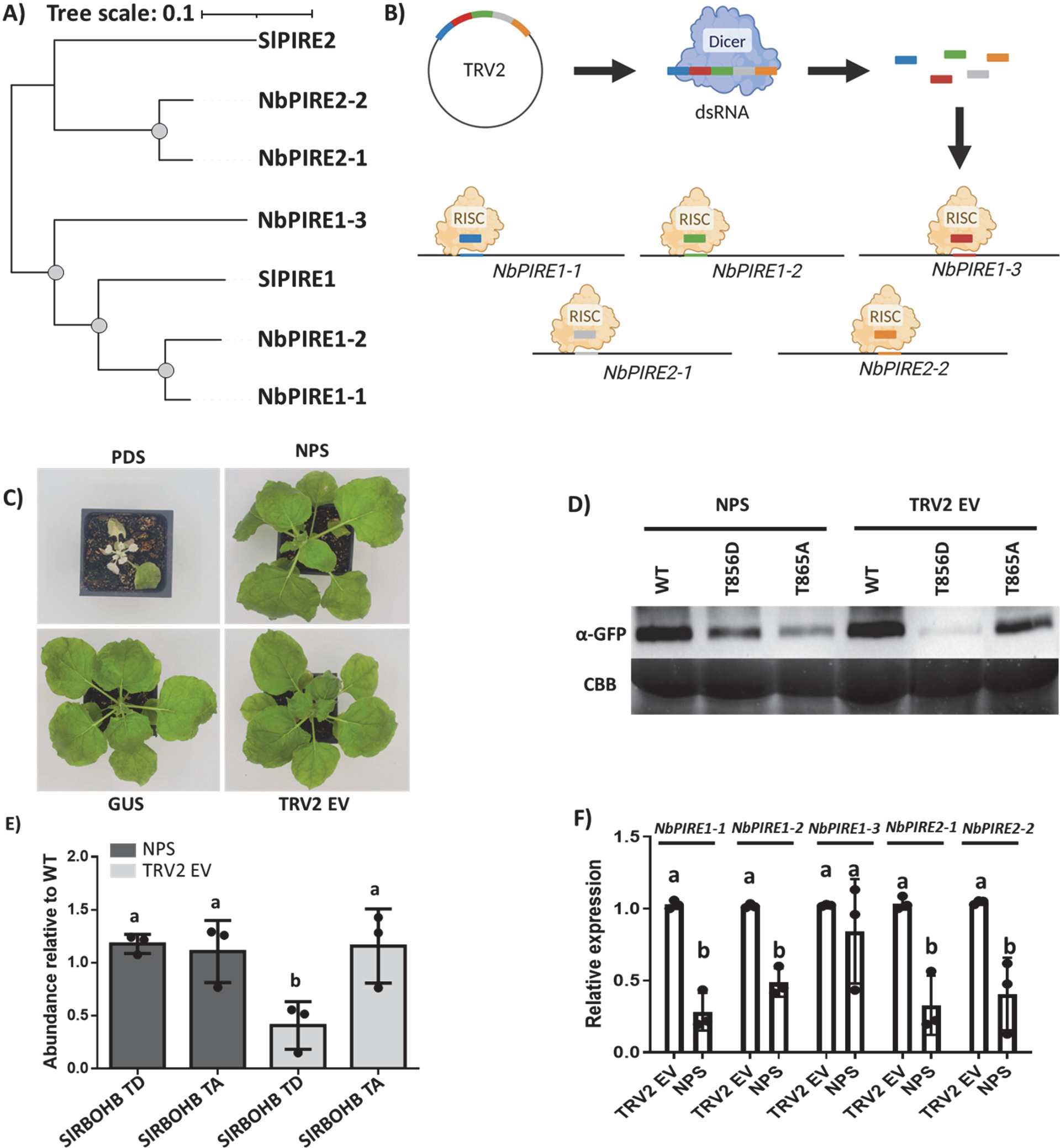
Changes in abundance of Phosphomimetic SlRBOHB is dependent on *PIRE* homologs. **A)** Phylogenetic tree of *Nicotiana benthamiana* and *Solanum lycopersicum* PIRE homologs. Sequence alignments were generated utilizing the clustal omega program and the mid-rooted phylogenetic tree was generated using maximum likelihood method with a bootstrap value of 1000. Grey dots signify bootstraps higher than 80, Sl = *S. lycopersicum*, Nb = *N. benthamiana*. **B)** Diagram of the stacked VIGS approach. Small (∼150bp) regions of *NbPIRE* homologs were cloned into the TRV2 silencing vector, then *Agrobacterium* carrying TRV1 and TRV2 were co-infiltrated into two-week-old *N. benthamiana*. The silencing fragments are converted into a long double stranded RNA (dsRNA) which then get processed by dicer to generate short interfering RNAs (siRNAs) leading to depletion of four out of five *NbPIRE* homologs (NPS construct). **C)** Images of *N. benthamiana* two-weeks post TRV inoculation via *A. tumefaciens*. The plant silenced for Phytoene Desaturase (*PDS*) displayed photobleaching and dwarfism. **D-E)** Wild-type SlRBOHB and phosphorylation mutants were transiently expressed in *N. benthamiana t*wo-weeks post TRV inoculation. Protein accumulation was visualized by anti-GFP immunoblotting and quantified utilizing Image Lab. Protein levels were first normalized using the Coomassie brilliant blue (CBB) signal. The relative intensity of the proteins was compared to RBOHB^WT^. N=3 blots, error bars display standard deviation. Statistical differences were calculated by ANOVA with post-hoc Tukey test (p-value = 0.0175). Silencing of *NbPIRE* homologs leads to enhanced accumulation for RBOHB^T856D^. TRV2^NPS^ plant displayed enhanced accumulation of RBOHB^T856D^ when compared to the TRV2 EV silencing control. **F)** *N. benthamiana* silenced plants and controls were subjected to qPCR to analyze *PIRE* expression levels. Relative expression was calculated compared to the Ef1α housekeeping gene. TRV2^NPS^ treated plants displayed significantly lower expression levels of *NbPIRE* homologs when compared to TRV2^EV^ control, except for *NbPIRE1-3*. Each data point represents the average of one biological replicate (N=3 plants), error bars = SD. Differences were detected by two-way ANOVA with a Sidak’s multiple comparison test (p value < 0.0001).

After identifying these *NbPIRE* homologs, virus induced gene silencing (VIGS) was performed to ascertain their role in SlRBOHB abundance. We used tobacco rattle virus (TRV), which replicates via a double-stranded RNA intermediate, for VIGS in *N. benthamiana* (Bekele et al., 2019; Rössner et al., 2022; Senthil-Kumar & Mysore, 2011). We simultaneously attempted to silence all *PIRE* homologs using a stacked VIGS approach which incorporates small (150bp) regions in a single TRV2 construct to silence *NbPire* homologs in parallel (TRV2^NPS^, NPS = *Nicotiana PIRE* Silencing) (**Fig. 3B**) (Ahn et al., 2023). To ensure *PIRE* silencing in *N. benthamiana,* we performed qPCR analysis. When compared to the TRV2^EV^ control, TRV2^NPS^ displayed significantly lower expression for all homologs except *NbPIRE 1-3*. (**Fig. 3F**).

To test the abundance of SlRBOHB phosphomutants after silencing *NbPIRE*s, we infiltrated *N. benthamiana* plants with TRV2^NPS^, TRV2^Gus^, TRV2^EV^ and TRV2^PDS^. As a control we used TRV2^PDS^ to silence phytoene desaturase (PDS) which interferes with the carotenoid biosynthesis pathway and induces photobleaching (Senthil-Kumar & Mysore, 2011) (**Fig. 3C**). Accumulation of SlRBOHB variants was assessed in silenced plants after *Agrobacterium-*mediated transient expression. Importantly, when multiple *PIRE* genes were silenced in *N. benthamiana*, the proteins SlRBOHB^WT^, SlRBOHB^T856A^, and SlRBOHB^T856D^ all exhibited similar accumulation levels (**Fig. 3D, E**). In contrast, plants treated with TRV2^EV^ displayed significantly lower levels of protein accumulation for SlRBOHB^T856D^ and similar protein accumulation for SlRBOHB^WT^ and SlRBOHB^T856A^ (**Fig. 3D, E**). These trends in protein accumulation were similar in TRV2^GUS^ treated plants (**Supplemental Fig. S3**) These results indicate changes in accumulation of the phosphomimetic SlRBOHB variants depend on *NbPIRE* in *N. benthamiana*.

### Gene editing of *SlPIRE1* leads to increased ROS production after flg22 treatment

In Arabidopsis, *pire* knockouts exhibit enhanced ROS production and disease resistance (Lee et al., 2020). Therefore, we hypothesize that targeting *PIRE* homologs in other plants may confer enhanced ROS production. CRIPR/Cas9 gene editing was used to generate *SlPIRE1* (Solyc03g113700) and *SlPIRE2* (Solyc06g071270) mutants. The CRISPR-P 2.0 web tool was used to select guide RNAs specifically targeting the N-terminal region of each homolog (**Fig. 4A, Supplementary Fig. S4**). Using these guide RNAs, we generated three different constructs to transform *S. lycopersicum* cv. M82. Two constructs targeted the *SlPIRE1* and *SlPIRE2* genes independently, while the third construct simultaneously targeted both genes. Two homozygous independent gene-edited lines for *SlPIRE1* (*Slpire1-1* and *Slpire1-2*) and *SlPIRE2* (*Slpire2-1* and *Slpire2-2*) (**Fig. 4A**) were obtained, which were verified by Sanger sequencing (**Supplementary Fig. S5**). These gene editing events led to the generation of frame-shift mutations leading to early stop codons. Cas9 was segregated out before conducting experiments with each line. After four rounds of transformation, and 40 independent transformants, we were unsuccessful in generating a double mutant, indicating it may be lethal.

**Figure 4.**
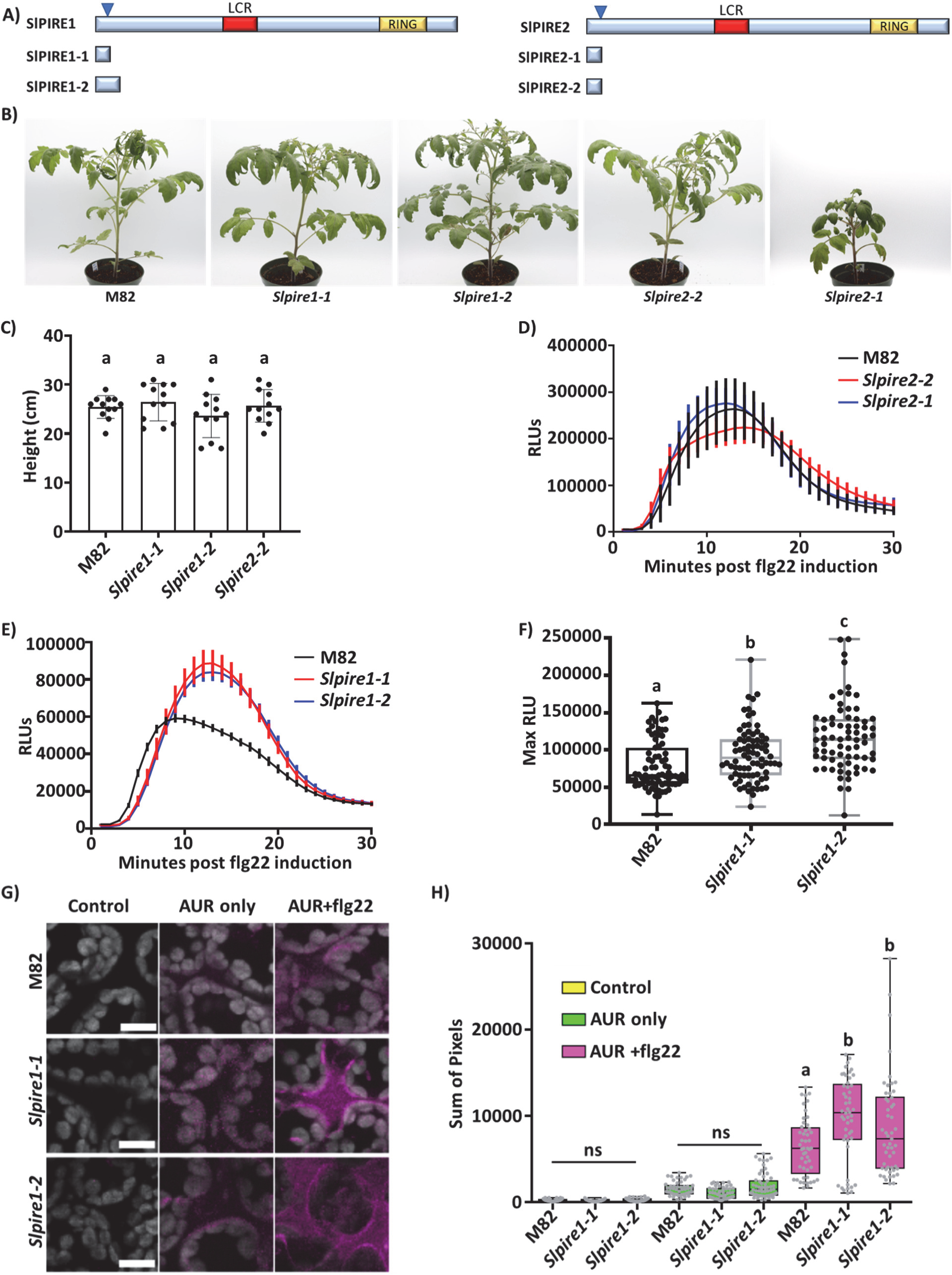
Editing tomato *SlPIRE1* results in enhanced production of reactive oxygen species upon flagellin perception. **A)** Diagram of SlPIRE1 and SlPIRE2, arrows represent areas targeted by CRISPR/Cas9. Below the protein diagrams are the predicted truncated proteins generated from gene editing in *Solanum lycopersicum* cv M82. **B)** The *SlPIRE1* gene edited lines did not display growth phenotypes in comparison to M82 (WT) plants, under vegetative growth conditions. The *Slpire2* line 1 (*Slpire2-1*) displayed decreased growth compared to M82, but *Slpire2-2* displayed growth rates similar to M82. **C)** Height quantification of M82 and gene edited lines. Heights were measured from soil to the shoot apical meristem. N= 15 plants. Statistical analysis was performed by ANOVA with post-hoc Tukey test (p-value = 0.2599) **(D-E)** ROS production was analyzed in four-week-old M82, *Slpire1,* and *Slpire2* after treatment with 100nM flg22. *Slpire2* gene edited lines did not display changes in ROS production in comparison to M82 after flg22 treatment. *Slpire1* lines displayed enhanced ROS production post flg22 treatment compared to M82. N = 3 plants with 8 leaf disks per plant, error bars = SEM, **F)** Quantification of ROS production in four-week-old M82, *Slpire1,* and *Slpire2* after treatment with 100nM flg22. Results display maximum relative light units (max RLU). *Slpire1* lines produce significantly higher max RLU compared to M82 after flg22 treatment. N=72 leaf disks over 3 sets of biological replicates (9 plants per genotype). Outliers were identified and removed using ROUT method (Q=1%). Statistical differences were calculated by a one-way ANOVA with post-hoc Tukey test (p value < 0.0001) **G-H)** ROS was visualized and quantified using the non-permeable Amplex Ultra Red (AUR) stain 15 minutes post-leaf infiltration with 100nM flg22. AUR was visualized by confocal microscopy. Representative images of M82, *Slpire1-1* and *Slpire1-2* with or without AUR and flg22 treatment. Image J was used to quantify the same size (1cm x1cm) of five randomly selected regions per image. Three plants per genotype with two images per leaf were quantified, n = 6 images per genotype and treatment. Outliers were identified and removed using ROUT method (Q=1%) differences were calculated by a one-way ANOVA with post-hoc Tukey test (p value < 0.0001). *Slpire1* lines exhibited significantly enhanced production of apoplastic ROS after induction with flg22.

The gene edited *Slpire1-1, Slpire1-2,* and *Slpire2-2* mutant lines displayed normal growth phenotypes in comparison to wild-type M82 plants (**Fig. 4B, C**). However, *Slpire2-1* exhibited low germination, delayed germination, and smaller stature compared to wild-type M82 (**Fig. 4B, C**). We analyzed flg22-induced ROS production on wild-type and gene edited lines. Both *Slpire2* lines and M82 produced similar levels of ROS after flg22 treatment (**Fig. 4D**). In contrast, *Slpire1-1* and *Slpire1-2* produced enhanced ROS compared to wild-type after flg22 treatment (**Fig 4E, F**). Therefore, we focused on *Slpire1* lines for future experiments. We also utilized Amplex UltraRed reagent (AUR), a membrane impermeable reagent that directly interacts with H_2_O_2_ to quantify ROS levels (Ashtamker et al., 2007; Cohn et al., 2008). Both *Slpire1-1* and *Slpire1-2* displayed significantly enhanced apoplastic ROS accumulation after flg22 induction (**Fig. 4G, H**). However, the baseline level of ROS in *Slpire1* edited lines and the dynamics of ROS production after flg22 treatment are not different from wild-type M82. Taken together, our data show that *Slpire1* gene edited lines specifically enhance apoplastic ROS production upon immune activation.

### *Slpire1* gene edited lines exhibit increased resistance to foliar bacterial pathogens

Since our gene edited *Slpire1* lines displayed enhanced production of ROS upon immune activation, we sought to test their ability to resist pathogen infection. We performed syringe infiltration of five-week-old plants with the bacterial strain *Pseudomonas syringae* pv. tomato DC3000 Δ*avrPto*Δ*avrPtoB* (DC3000ΔΔ), which contains mutations in two effectors and is less virulent than DC3000 (Lin & Martin, 2005). Both *Slpire1-1* and *Slpire1-2* exhibited reduced disease symptoms compared to wild-type M82 three days post-infection (**Fig. 5A**). Furthermore, *Slpire1* lines displayed an 18-fold reduction in bacterial titers compared to the M82 control (**Fig. 5B**). Since DC3000ΔΔ exhibits attenuated virulence, we also utilized wild-type *P. syringae* DC3000 to challenge *Slpire1* lines in the M82 background. Similar to the infections for DC3000ΔΔ, infections with DC3000 led to decreased disease symptoms and 15-fold decrease in bacterial accumulation (**Fig 5A,C**). Next, we investigated the role of *SlPIRE1* in disease resistance to the causal agent of bacterial spot of tomato, *Xanthomonas campestris* pv. *vesicatoria* (XCV 85-10). Infections with *X. campestris* led to decreased disease symptoms, including reduced chlorosis in *Slpire1* lines compared to M82 at seven days post-infection (**Fig. 5A)**. Bacterial titers for *X. campestris* were 12-fold lower for both *Slpire1* lines when compared to the wild-type M82 control (**Fig. 5D).** Collectively, these data demonstrate that *SlPIRE1* mutants exhibit higher defense induced ROS and increased disease resistance to foliar bacterial pathogens.

**Figure 5.**
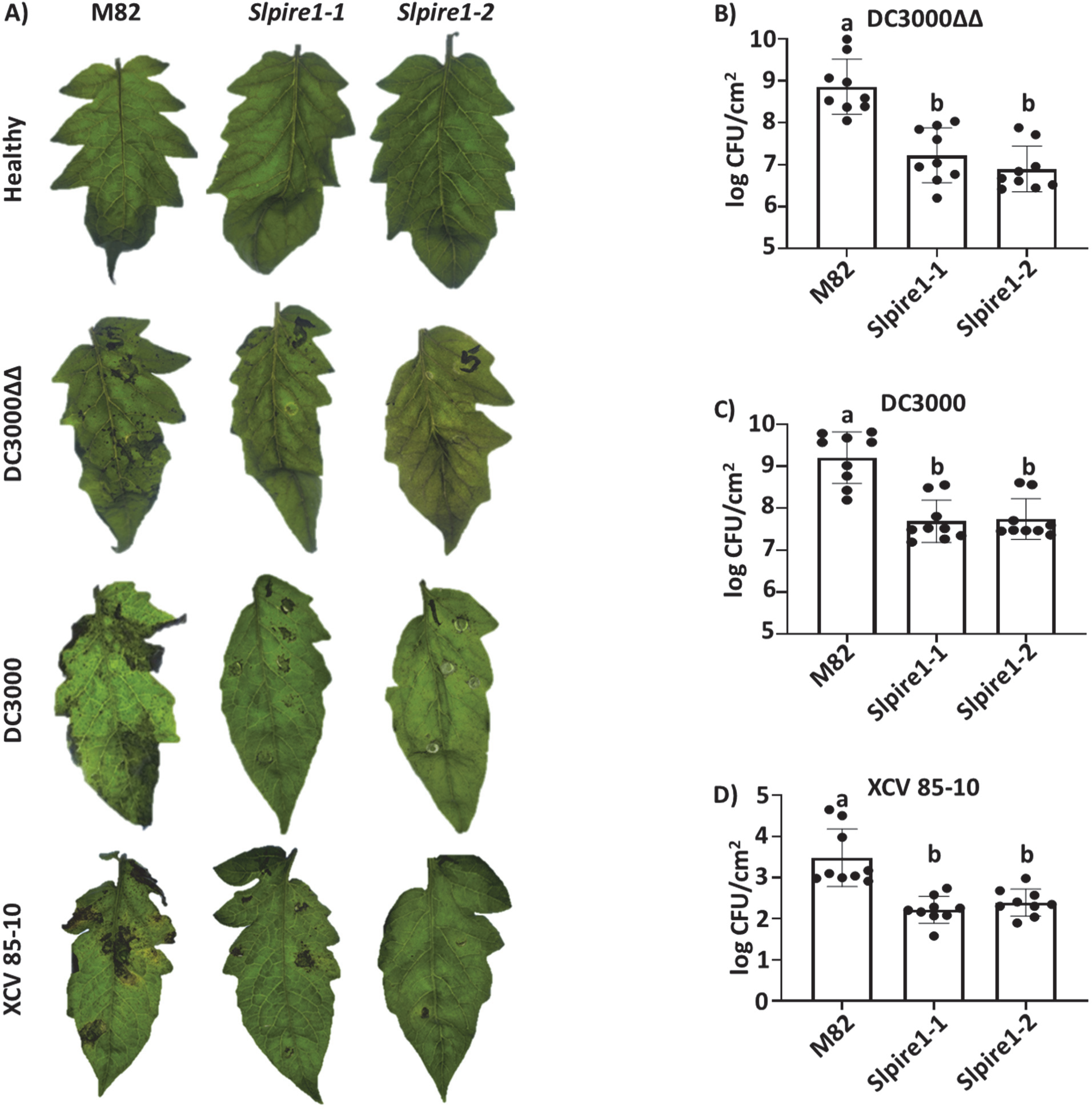
Editing *SlPIRE1* results in decreased disease symptoms and bacterial accumulation. **A)** Two independently gene edited *SlPIRE1* lines (*Slpire1-1* and *Slpire1-2*) displayed reduced disease symptoms 3 days post inoculation (dpi) with *Pst* DC3000 Δ*avrPto*Δ*avrPtoB* (DC3000ΔΔ), 3dp for *Pst* DC3000 (DC3000) and 7dpi for *Xanthomonas campestris* pv. *vesicatoria* (XCV 85-10). Representative images of 9 plant infections. **B-D)** To determine bacterial titers, leaf tissue was sampled 3dpi for DC3000 and 7dpi for XCV 85-10. Both *Slpire1-1* and *Slpire1-2* lines displayed decreased accumulation of DC3000ΔΔ **(B)**, DC3000 **(C)** and XCV 85-10 **(D)** compared to wild-type M82. n= 9 plants. Statistical analysis was performed by one-way ANOVA with post-hoc Tukey test (DC3000ΔΔ p-value < 0.0001, DC3000 p-value < 0.0001, and XCV85-10 p-value = 0.0423)

### *Slpire1* gene edited lines do not impact disease caused by root colonizing pathogens

To more comprehensively understand the role of *SlPIRE1* in plant defense, we also investigated its impact on root-invading pathogens. First, we challenged our gene edited lines with *Ralstonia pseudosolanacearum* GMI1000, a soil-borne Gram-negative bacteria that causes bacterial wilt by colonization of xylem vessels (Ingel et al., 2022; Lowe-Power et al., 2016; Salanoubat et al., 2002). After petiole inoculation, plants were monitored for 14 days, and the disease index was measured. There were no significant differences in the disease index between wild type M82 and *Slpire1-1* or *Slpire1-2* (**Supplemental Fig. S6A**). We also did not detect a difference after soil drench with *R. solanacearum* (**Supplemental Fig. S6B**). Next, we challenged *Slpire1-1* against the root-knot nematode *Meloidogyne javanica. M. javanica* invades the root tip and travels to the vascular cylinder to establish feeding sites comprised of giant cells. Here the root-knot nematode will remain sedentary and complete its life cycle (Bartlem et al., 2014). We assessed nematode infection seven weeks post inoculation by extracting and quantifying *M. javanica* eggs from infected roots as a proxy for disease progress. We did not detect differences in egg accumulation between wild-type M82 and *Slpire1-1* (**Supplemental Fig. S6C**). Taken together, these data indicate that targeting *SlPIRE1* enhances foliar disease resistance without affecting root-colonizing pathogens.

## Discussion

For decades *Arabidopsis thaliana* has been utilized as an effective model system to study plant immunity. Arabidopsis is favored for its short life cycle, and the extensive tools available for genetic manipulation (Nishimura & Dangl, 2010; Rédei, 1975). The discovery and investigation of Arabidopsis NLR and PRR immune receptors have provided insight into how plants recognize pathogens and activate immunity (Bent et al., 1994; Jones et al., 2024; Zipfel et al., 2006). Our knowledge of downstream immune signaling components stems from work in conducted Arabidopsis (Couto & Zipfel, 2016; Jones et al., 2024; Yuan et al., 2023). These findings have laid the foundation for potential translation to crop plants. For example, the Arabidopsis PRR Elongation Factor Tu Receptor (EFR) has been successfully introduced to multiple crop species, including tomato, rice, and sweet orange resulting in resistance to a variety of bacterial pathogens (Kunwar et al., 2018; Lu et al., 2015; Mitre et al., 2021). FLS2^XL^, a homolog of Arabidopsis FLS2, from wild grape can recognize *Agrobacterium,* a pathogen with divergent flg22 epitopes (Fürst et al., 2020). In this study, we examined the importance of the E3 ligase *PIRE*, which was originally identified in Arabidopsis, for its function in *S*. *lycopersicum.* By targeting *PIRE* homologs, we modulated the abundance of RBOHs in solanaceous plants and increased disease resistance to both *P. syringae* and *X. campestris*. This study highlights another immune regulator originally identified in Arabidopsis with promise to enhance disease resistance in a variety of plant species.

The versatility of CRISPR/Cas9 to target genes across multiple plant systems has been leveraged to target susceptibility (S) genes for disease control (Bisht et al., 2019; van Schie & Takken, 2014). Different classes of S genes include those involved in pathogen penetration, negative regulation of immune responses, and pathogen proliferation/dissemination (van Schie & Takken, 2014). *PIRE* is a negative regulator of immune responses and regulates RBOH stability and ROS production in Arabidopsis (Lee et al., 2020). Our results indicate *SlPIRE1* is a promising S gene that also negatively regulates immune responses and foliar pathogen accumulation in tomato. There have been multiple examples of gene editing of negative immune regulators leading to increased disease resistance. Recently, gene editing of the xylem sap protein 10 (*XSP10*) and salicylic acid methyl transferase (*SISAMT*) led to tolerance to Fusarium wilt disease in tomato (Debbarma et al., 2023). Another well-known example is the negative immune regulator, Mildew Locus O (*MLO)*. *MLO* mutants exhibit enhanced resistance to powdery mildew fungi in barley, wheat, and tomato (Jacott et al., 2021). However, production of higher order *mlo* mutants result in negative growth/yield penalties, including premature leaf senescence (Acevedo-Garcia et al., 2017; Jacott et al., 2021). Recently, the pleiotropic effects of *mlo* in *Triticum aestivum* have been circumvented by generating targeted mutations which lead to enhanced transcription *TaTMTB3,* a gene located directly upstream of *Mlo* on the chromosome, which uncouples negative growth phenotypes and resistance (Li et al., 2022). The most ideal S genes are those like *SlPIRE1,* where resistance is uncoupled from other pleiotropic effects. However, further characterization is necessary to ensure *SlPIRE1* lines do not display altered growth or yield phenotypes under field conditions.

*Slpire1* gene edited lines did not display higher baseline apoplastic ROS but generated enhanced ROS production upon PRR activation. It is likely that higher baseline levels of SlRBOHB result in increased ROS production upon pathogen perception. In Arabidopsis, ROS production by RBOHD requires activation via calcium binding and phosphorylation (Kadota et al., 2015; Li et al., 2014; Thor et al., 2020; Tian et al., 2019; Zhang et al., 2018). Phosphorylation of Arabidopsis RBOHD at T912 leads to PIRE-mediated ubiquitination and vacuolar degradation, regulating the level of steady-state RBOHD (Lee et al., 2020). In our experiments, we observed significantly higher ROS production in *Slpire1* after induction with flg22, but not at a resting state (**Fig. 4E-H**). This is consistent with the requirement of RBOHs to be post-translationally modified upon pathogen perception to generate ROS. Here we show that mutations of the corresponding T912 residues in SlRBOHB (**Fig. 2D, E**) or mutations in *SlPIRE1* (**Fig. 3D-F**) lead to changes in SlRBOHB accumulation. Taken together this suggests a model where SlRBOHB steady state accumulation is enhanced by removal of *SlPIRE1* which leads to increased ROS production upon pathogen perception.

E3 ligases are important as they provide specificity and bridge the interaction between the E2 ubiquitin ligase and their target protein (Sadanandom et al., 2012). Interestingly, neither of the *Slpire2* edited lines displayed alterations in defense-induced ROS production. This suggests that PIRE homologs in tomato do not have completely overlapping targets. Our inability to acquire the double mutant line for *Slpire1* and *Slpire2* suggests that removing both may be lethal. RBOHs play a role in plant development as well as response to stress (Kadota et al., 2015). In *Arabidopsis*, processes such as pollen tube growth, seed ripening, and formation of root hairs are dependent on *AtRBOHH, AtRBOHJ, AtRBOHB* and *AtRBOHC,* respectively (Kaya et al., 2014; Lassig et al., 2014; Müller et al., 2009; Takeda et al., 2008). It is possible that *Slpire1* and *Slpire2* collectively regulate other RBOHs in *S. lycopersicum*.

Although *SlPIRE1* acts as an S gene towards the foliar pathogens *Pseudomonas* and *Xanthomonas*, it does not affect disease development for the root colonizing bacteria *R. pseudosolanacearum* or root-knot nematode *M. javanica*. For pathogens with different life cycles, S genes can lead to enhanced susceptibility. For example, targeting the S gene *mlo* confers resistance to powdery mildew in wheat, but enhances susceptibility to *Magnaporthe oryzae* pathotype *Triticum* (Gruner et al., 2020). However, there is evidence that activation of plant immune receptors can restrict both root-colonizing pathogens we tested. The NLR *Mi-1* has been used for decades to control resistance to root-knot nematodes within the MIG group (*Meloidogyne incognita* group) and is incorporated into many commercial tomato cultivars (Wubie & Temesgen, 2019.). Transfer of the *EF-Tu* PRR to *S. lycopersicum* confers resistance to *R. pseudosolanacearum* in both greenhouse and field conditions (Kunwar et al., 2018; Lacombe et al., 2010). Targeting two enzymes involved in PRR-induced ROS, overexpression of the *RIPK* kinase or genome editing of the protein phosphatase *LOPP*, in the dwarf *S. lycopersicum* model plant, Micro-Tom, resulted in increased resistance to *R. pseudosolanacearum* (Wang et al., 2022). Roots are in contact with diverse microorganisms and ROS can induce proliferation and induction of lateral roots, which are sites of entry for both *R. pseudosolanacearum* and *M. javanica* (Hasan et al., 2024; Manzano et al., 2014; Tarkowski et al., 2023; Vailleau & Genin, 2023). It is possible that the inhibitory effect of increasing ROS in *Slpire1* is counteracted by alterations in root architecture. Alternatively, *SlPIRE1* may exhibit a different function or targets in root versus leaf tissue, consistent with the specificity achieved by RBOHs in different plant tissues (Chen & Yang, 2020).

Pathogens frequently overcome single gene resistance, and no single R or S gene can serve as a silver bullet against all pathogens. A multilayered strategy that integrates resistance mechanisms at different stages of infection is a promising approach for durable disease resistance (Zhang & Coaker, 2017). In *Oryza sativa*, expression of the PRR *Xa21* along with mutations of S genes including the transcription factor subunit *Xa5* and the sugar transporter *Xa13,* leads to resistance against *Xanthomonas oryzae* (Akter et al., 2024; Huang et al., 1997). Pyramiding a minimum of two adult plant resistance genes in *Triticum aestivum* resulted in adequate seedling stage resistance to stripe rust, caused by *Puccinia striiformis* f. sp. *tritici* (Wang et al., 2023). Targeting *PIRE,* in combination with other loci, could be a promising approach for effective pathogen control in the future.

## Materials and Methods

### Plant growth conditions

*Nicotiana benthamiana* was grown in a controlled environment chamber at 26°C with a 16-h light/8-h dark photoperiod (180 μmol m−2 s−1). Four-week-old plants were used for *Agrobacterium*-mediated transient protein expression. Tomato plants (*Solanum lycopersicum* cv. M82) were grown under controlled conditions at 26°C and 12-h light/12-h dark photoperiod. Five-week-old tomatoes were used for height measurements, ROS assays and pathogen challenge. For height measurements plants were measured from soil to the shoot apical meristem.

### Gene editing: guide design and construct generation

CRISPR guide RNAs (gRNAs) were designed using the CRISPR-P 2.0 web tool (http://crispr.hzau.edu.cn/CRISPR2). gRNAs were selected based on early targeting of the *SlPIRE1* and *SlPIRE2* genes, an on-target score higher than 0.4, and off-targets with scores primarily lower than 0.5 and in intergenic regions. gRNAs and off-target analysis are provided in **Supplemental Tables S2** and **S3**. For single gRNA constructs, gRNAs were cloned into the pCR3-EF plasmid containing Cas9 (Fister et al., 2018) using Golden Gate assembly utilizing the *Bsa*I-HFv2 (NEB E1601S) restriction enzyme.

For multiplex constructs targeting *SlPIRE1* and *SlPIRE2*, gRNA primers were used to amplify tRNA between both gRNAs (Xie et al., 2015). This gRNA-tRNA-gRNA multiplex was cloned into pCR3-EF plasmid using Golden Gate assembly as described above. pCR3-EF constructs containing the gRNAs were recombined into the pPZP200 destination vector (Hajdukiewicz et al., 1994). pPZP200 constructs containing gRNAs targeting *SlPire1, SlPire2* and *SlPire1/SlPire2* were transformed into the cultivar M82 via *Agrobacterium* at the Innovative Genomics Institute (IGI Berkeley) and the transformation facility at University of Nebraska-Lincoln Center for Plant Sciences. Gene edited lines were confirmed by Sanger sequencing for the targeted genes. Primers are listed in **Supplemental Table S4**.

### Sequence and phylogenetic analyses of PIRE homologs

Plant PIRE homologs were mined in NCBI utilizing BLASTP. We used mined homologs using the modified RING-C2 domain found in AtPIRE (AT3g48070). Utilizing this strategy, we identified 170 modified RING-C2 domain proteins with >70% amino acid (aa) similarity to the AtPIRE modified RING domain in Charyophyta, Bryophyta, Gymnosperms, and Angiosperms. Full-length proteins were aligned utilizing Clustal Omega. Phylogenetic trees based on the RING domain of identified PIRE homologs were generated using the maximum likelihood method with a bootstrap value of 1000 in IQ-TREE (Minh et al., 2020; Nguyen et al., 2015). Protein domains and low complexity regions were identified utilizing SMART (Simple Modular Architecture Research Tool) (Letunic et al., 2021). For *N. benthamiana* PIRE homologs we used the SlPIRE1 (Solyc03g113700) and SlPIRE2 (Solyc06g071270) aa sequence to mine for homologs. The NbPIRE1-1 (Niben101Scf04654g02005), NbPIRE1-2 (Niben101Scf07162g01018), NbPIRE1-3 (Niben101Scf07162g01018), NbPIRE2-1 (Niben101Scf02237g01001), and NbPIRE2-2 (Niben101Scf06720g01006) aa sequences were aligned using Clustal Omega. Phylogenetic trees were generated with the maximum likelihood method with a bootstrap value of 1000 in IQ-TREE. **Supplementary Table S1** includes the gene identifiers of all PIRE homologs.

### Transient expression in *Nicotiana benthamiana*

For transient expression experiments, we generated constructs of SlRBOHB (Solyc03g117980) with C-terminal fusions to YFP. PCR amplified cDNA was then directionally cloned into pENTR/D-TOPO (Invitrogen). Site-directed mutagenesis was performed on the pENTR/D-TOPO construct containing RBOHB was performed to generate phosphomutants. pENTR/D-TOPO constructs were then sequenced before recombination into the pEarleyGate104 destination vector by LR reaction (Earley et al., 2006). Constructs were electroporated into *Agrobacterium tumefaciens* (GV3101). Leaves of four-week-old *N. benthamiana* were infiltrated with the *Agrobacterium* containing each of the generated constructs (SlRBOHB^WT^, SlRBOHB^T856A^, SlRBOHB^T856D^, and EV) (OD_600_=0.6). Leaf tissue was harvested 48 hours post infiltration (hpi) (3 leaf disks #3 cork borer (7mm) per sample). Tissue was ground in 100µl of Laemmli buffer (Laemmli, 1970). Protein samples were separated by SDS-PAGE and immunoblotting was performed using anti-GFP-HRP at a concentration of 1:5000 (Miltenyi Biotec, 130-091-833, clone GG4-2C2.12.10). Image intensity quantifications were performed using Image Lab software (Image lab software version 6.1). All experiments were repeated at least three times with similar results. Data were analyzed by a Kruskal-Wallis test with a Dunn’s test (p-value: 0.0003).

### ROS burst assay

In *N. benthamiana,* leaf disks (4mm diameter) were collected from plants transiently expressing SlRBOHB^WT^, SlRBOHB^T856A^, SlRBOHB^T856D^, and EV on the same leaf. Leaf disks were placed in water (200 µl) for 20 hrs in CorningTM CorstarTM 96-well solid plates (Fisher #07-200-589) to recover before inducing with flg22. ROS was measured as previously described (Lee et al., 2020). The reaction solution contained 20 μM L-012 (a luminol derivative from Wako Chemicals USA #120-04891), 10 mg mL−1 horseradish peroxidase (Sigma), and 100nM flg22 (GeneScript, 95% purity). Light intensity was measured using a TriStar LB 941 plate reader (Berthold Technologies). In *S. lycopercicum*, eight leaf disks (4mm) were collected per plant per genotype (M82, *Slpire1-1*, *Slpire1-2, Slpire2-1, Slpire2-2*). The assay was performed as described above. All experiments were repeated at least three times with similar results. Data from three experiments were combined. Whiskers show minimum and maximum values. Statistical differences were determined by ANOVA with post-hoc Tukey test (p-value: 0.0001).

### Virus induced gene silencing (VIGS) of *NbPIRE* homologs

For VIGS a gene block was generated (Twist Bioscience) containing 150 bp long regions of each *NbPIRE* homolog, cloned into pENTR/D-TOPO (Invitrogen), and recombined into TRV2 destination vector via LR clonase reaction. The TRV2 construct along with the TRV1 constructs were then electroporated into *Agrobacterium* (GV3101). Two-week-old *N. benthamiana* plants were co-infiltrated (OD600 = 0.4) with *Agrobacterium* containing TRV1 and TRV2 with one specific silencing region (TRV2^NPS^, TRV2^GUS^, TRV2^EV^ and TRV2^PDS^). *N. benthamiana* plants were allowed to grow for another two weeks after infiltrations (four-week-old plants) before transient expression. TRV2^PDS^ VIGS (Xu et al., 2019) plants served as a control to monitor silencing progress. Transient expression was performed as described above. Briefly *N. benthamiana* leaves were infiltrated with *Agrobacterium* containing the SlRBOHB variants described above. Leaf tissue was harvested 48hpi and ground in 100µl of Laemmli buffer (Laemmli, 1970). Protein samples were separated via SDS-PAGE gel and immunoblotting was performed using anti-GFP-HRP at a concentration of 1:5000 (Miltenyi Biotec, 130-091-833, clone GG4-2C2.12.10). Image intensity quantifications were performed using Image Lab software (Image lab software version 6.1). All experiments were repeated at least three times with similar results. Data were analyzed by ANOVA with post-hoc Tukey test (alpha = 0.05).

### qPCR of VIGS silenced plants

To examine the expression of *NbPire* homologs after silencing, we harvested three-leaf punches with a #3 cork borer (7 mm) at the same time as we collected tissue for transient expression. Tissue was frozen and ground using liquid nitrogen. RNA was extracted from these plant samples with TRIzol (Fisher #15596018), following the manufacturer’s instructions. DNase treatments for RNA preps were performed with RQ1 RNase-Free DNase (Promega #PR-M6101). cDNA synthesis was performed with the MMLV Reverse Transcriptase (Promega #PRM1705) kit. Primers for qPCR were designed using Primer3 (Untergasser et al., 2012) and are found in Supplemental Table S4. Gene expression was calculated using the Ct method and was normalized against the *N. benthamiana* EF1a housekeeping gene. qPCR reactions were performed with SsoFast EvaGreen Supermix with Low ROX (BioRad #1725211) in a 96-well white PCR plate (BioRad #HSP9601) according to the manufacturer’s instructions. All experiments were repeated at least three times with similar results. Graphed data represent three biological replicates and differences were detected by two-way ANOVA (alpha = 0.05).

### Visualization of apoplastic ROS by AUR

To visualize apoplastic ROS, five-week-old *S. lycopersicum* plant leaves were syringe infiltrated with Amplex Ultra Red (AUR) or a combination of AUR with 100nm flg22 (GeneScript, 95% purity). Leaf tissue was then visualized by confocal microscope (Leica TCS SP8) 15 minutes post infiltration. Control images were taken from tissue that was not infiltrated. Images were taken from five randomly selected regions of the same size (1cm x1cm). Images were analyzed through ImageJ. Threshold: Default mode and minimum 15 intensity were used for all images. RawintDen (the sum of the values of the pixels in the image or selection) was used to quantify and compare. Three plants per genotype, two imaged per plant, and a total of six images for each treatment were quantified. Outliers were identified and removed using ROUT method (Q=1%) differences per treatment were calculated a one-way ANOVA with post-hoc Tukey test (alpha = 0.05).

### Disease assays

The *Pseudomonas syringae* pv. *tomato* DC3000 (DC3000), *Pseudomonas syringae* pv. *tomato* DC3000 Δ*avrPto*Δ*avrPtoB* (DC3000ΔΔ) and *Xanthomonas campestris* pv. *vesicatoria* (XCV 85-10), were grown on NYG plates (Liu et al., 2013) with the appropriate antibiotics two days prior to infiltration. On the day of infection DC3000ΔΔ, DC3000 and XCV85-10 were resuspended on 5mM MgCl_2_ (DC3000ΔΔ and DC3000 OD600 = 0.00005, XCV85-10 OD600 = 0.0003). Five-week-old M82, *Slpire1-1*, and *Slpire1-2* plants were syringe inoculated with the pathogens listed above. We inoculated three to four leaves per plant per genotype (n=5 plants per experiment). Leaf tissue was collected 3 days post inoculation (dpi). Images were collected from representative leaves. To measure the bacterial titers, we collected one #3 (7 mm) leaf disk per infected tissue. Leaf disks were ground up in 200μl of 5mM MgCl_2_ and the solution was serially diluted from 10^−1^ to 10^−7^. Serial dilutions were plated on NGY plates containing appropriate antibiotics along with 50μg/ml of cycloheximide. Colony counts were then performed after incubation at 28°C for 48 h to determine the log CFU/cm^2^. All experiments were repeated at least three times with similar results. Statistical analysis was done by one-way ANOVA with post-hoc Tukey test (DC3000ΔΔ p value: 0.0001, DC3000 p value: 0.0001, and XCV85-10 p value: 0.0423)

For infection with *R. pseudosolanacearum* GMI1000, we inoculated 21-day-old tomato plants with a cut-petiole approach by excising the lowest petiole and inoculating its surface with a 2-µl droplet of 5×10^5^ cfu/mL bacterial suspension (Khokhani et al., 2018). We rated disease progress for 14 days following disease index scale from 0 to 4, where 0 = 0 wilted leaves, 1 = 0.1 to 25% of wilted leaves, 2 = 25.1 to 50% of wilted leaves, 3 = 50.1 to 75% of leaflets wilted, and 4 = 75.1 to 100% of wilted leaves (Khokhani et al., 2018)

For *M. javanica* infections, sterile nematode eggs were collected from previously infected tomato plant cultures (MV variety) using a 10% bleach solution. Eggs were allowed to hatch at 27°C to collect J2 stage nematodes. Collected J2 stage nematodes where then washed on a 50 mL vacuum filtration unit (e.g., 22 µm, Thermo Scientific Nalgen Filtration Product, USA) using sterile water. J2 numbers were obtained at this point before infecting 4-week-old plants (M82 and Slpire1-1). Infected plants were harvested 7 weeks post infection. Eggs were collected from infected plants by using a 10% bleach wash and using sieves of mesh #200 (75 µm) and mesh #500 (25 µm) to separate the eggs. Egg counts were performed under a dissecting microscope. All experiments were repeated twice with similar results, data were analyzed for significant differences by t-test.

### Data Availability

All plasmids will be deposited in Addgene upon acceptance. All raw data have been deposited in Zenodo: https://doi.org/10.5281/zenodo.13119655.

## Supplemental Data

**Supplementary Dataset S1.** Identified PIRE homologs.

**Supplementary Table S1.** *SlPIRE1* gRNA analysis

**Supplementary Table S2.** *SlPIRE2* gRNA analysis

**Supplementary Table S3.** qPCR primers

**Supplementary Figure S1.** Alignment of the modified RING-C2 domain found in Green Algae, Bryophytes, Gymnosperms, and Angiosperms

**Supplemental Figure S2.** Transient expression of SlRBOHB can be differentiated from the endogenous NbRBOHB burst after flg22 induction.

**Supplemental Figure S3.** SlRBOHB phosphomutant expression is similar for the GUS and TRV2 controls in *Nicotiana benthamiana*.

**Supplemental Figure S4.** Validation of *Slpire* gene edited lines by DNA sequencing.

**Supplemental Figure S5.** Alignments of the amino acid translations for gene edited *Slpire* lines.

**Supplemental Figure S6.** Disease measurements for *R. pseudosolanacearum* GMI1000 strains and *M. javanica* strains on M82 (wild-type) and *Slpire1* edited lines.

## Author Contributions and Acknowledgements

GC and BC designed the research, analyzed the data, and wrote the paper. BC performed most experiments under the guidance of GC unless otherwise noted. SB and MT assisted with *Pseudomonas* and *Xanthomonas* infections and genotyped edited lines. AT genotyped edited lines. JZ performed AUR staining and image analyses. TL performed phylogenetic analyses. NA performed *Ralstonia* infections. AB and BC performed *Melodogyne* infections. MG, EZ, and M-JC generated edited lines. TL-P, SS, BS and M-JC helped design the research for pathogen infection and genome editing.

