## Supplementary Tables S1-S3, Figures S1-S6 for "Gene editing of the E3 ligase *PIRE1* fine-tunes ROS production for enhanced bacterial disease resistance in tomato"

**Supplementary materials:**

**Dataset S1, Tables S2-S3, Figures S1-S6**

**Supplemental Tables**

**Supplemental Table S1. *SlPIRE1* gRNA analysis**

Table displays the position of the gRNA, gRNA sequence, and off-target sites with scores.

*Slpire1* guide

| | | | | | |
|---|---|---|---|---|---|
| guide1 on-score: 0.4078 | | | | | |
| postion: SL3.0ch03:-65252750 | | | | | |
| guide sequence: TTGGGGTTAGAGGAAGCAATAGG | | | | | |
| number of offtarget sites: 91 | | | | | |
| top 20 genome-wide off-target sites | | | | | |
| Sequence | Off-score | MMs | Locus | Gene | Region |
| TTGGGATTAAAGGAAGCAATGGG | 0.933 | 2MMs | SL3.0ch04:-39722100 | gene:Solyc04g049130.3 | intron |
| TTAGAGTTAGAGAAAACAATAGG | 0.6 | 4MMs | SL3.0ch08:-48082971 | | Intergenic |
| TTGGGCTGAGAAGAAACAATAGG | 0.467 | 4MMs | SL3.0ch02:-15727873 | gene:Solyc02g014170.3 | intron |
| TTGGGGCGAGAAAAAGCAATAGG | 0.434 | 4MMs | SL3.0ch02:-32851223 | | Intergenic |
| TTGGGGAAAGAGGAAGAAATAGG | 0.327 | 3MMs | SL3.0ch03:-30995822 | gene:Solyc03g059316.1 | intron |
| TTGGGGAAAAAGGAAGAAATGGG | 0.305 | 4MMs | SL3.0ch09:-14872449 | | Intergenic |
| TTGGGGAAAAAGGAAGAAATGGG | 0.305 | 4MMs | SL3.0ch04:-13773084 | | Intergenic |
| TTGGGGAAAAAGGAAGAAATGGG | 0.305 | 4MMs | SL3.0ch09:-46235736 | | Intergenic |
| TTGGGGAAAAAGGAAGAAATGGG | 0.305 | 4MMs | SL3.0ch10:+28569421 | | Intergenic |
| TTGGGGAAAAAGGAAGAAATGGG | 0.305 | 4MMs | SL3.0ch08:+12339840 | | Intergenic |
| TTGGGGAAAAAGGAAGAAATGGG | 0.305 | 4MMs | SL3.0ch09:+48321502 | | Intergenic |
| TTGGGGTGAAAGAAAGAAATGGG | 0.295 | 4MMs | SL3.0ch01:+55850960 | | Intergenic |
| TTGAGGTTAAGGGAAGCATTGGG | 0.294 | 4MMs | SL3.0ch08:-316739 | gene:Solyc08g005430.3 | utr |
| TTGGGGCTAGAACAAGCAATAGG | 0.27 | 3MMs | SL3.0ch09:+66041209 | | Intergenic |
| TTGGAGGAAGAGGAGGCAATGGG | 0.265 | 4MMs | SL3.0ch09:-7380213 | gene:Solyc09g014980.3 | CDS |
| TTGGGATTTGAACAAGCAATAGG | 0.236 | 4MMs | SL3.0ch09:+41852222 | | Intergenic |
| TGGGGGTATGAGGAAGCAAATGG | 0.227 | 4MMs | SL3.0ch03:-64463534 | gene:Solyc03g112590.3 | CDS |
| TTGGGGTTAGAAAAAGAAAAAGG | 0.226 | 4MMs | SL3.0ch03:-38929962 | | Intergenic |
| TTTAGGGAAGAGGAAGCAATTGG | 0.212 | 4MMs | SL3.0ch03:+10672548 | | Intergenic |
| TTGGGGGAAAAGGAAGAAATAGG | 0.205 | 4MMs | SL3.0ch06:+5875458 | | Intergenic |

**Supplemental Table S2. *SlPIRE2* gRNA analysis**

Table displays the position of the gRNA, gRNA sequence, and off-target sites with scores.

### *Slpire2* guide

| | | | | | |
|---|---|---|---|---|---|
| guide1 on-score: 0.4317 | | | | | |
| postion: SL3.0ch06:-43989211 | | | | | |
| guide sequence: TGGAATTAGAGGAAGCGACAGGG | | | | | |
| number of offtarget sites: 29 | | | | | |
| top 20 genome-wide off-target sites | | | | | |
| Sequence | Off-score | MMs | Locus | Gene | Region |
| TGGAACTATAAGAAGCAACAAGG | 0.453 | 4MMs | SL3.0ch02:-54049947 | gene:Solyc02g092290.3 | CDS |
| TGGAATTATTGGAAGCAACATGG | 0.439 | 3MMs | SL3.0ch02:+39993027 | gene:Solyc02g069580.3<br>gene:Solyc02g069570.3 | CDS |
| TGGAATTTGAAGAAGAAACAGGG | 0.4 | 4MMs | SL3.0ch03:+2095648 | | Intergenic |
| TTGAATGAGAGGAAGAAACAGGG | 0.349 | 4MMs | SL3.0ch11:-47658739 | | Intergenic |
| TTGAATGAGAGGAAGAAACAGGG | 0.349 | 4MMs | SL3.0ch11:-47655215 | | Intergenic |
| TGGAAATTGAAGAAGCAACAGGG | 0.347 | 4MMs | SL3.0ch07:-815344 | gene:Solyc07g005970.3 | intron |
| TGGAAGAAGAGGAAGAGACTAGG | 0.3 | 4MMs | SL3.0ch03:+46056661 | | Intergenic |
| TGGAGATAGAGGAGGAGACAAGG | 0.291 | 4MMs | SL3.0ch05:+57118290 | | Intergenic |
| TGGGATTTGAGGAAAAGACATGG | 0.252 | 4MMs | SL3.0ch07:+54493200 | | Intergenic |
| TGGGGTTAGAGGAAGCAATAGGG | 0.194 | 4MMs | SL3.0ch03:-65252755 | gene:Solyc03g113700.3 | CDS |
| AGGAAGTGGAGGAAGCGAAATGG | 0.18 | 4MMs | SL3.0ch04:-20397120 | | Intergenic |
| TAGTATTAGAGGAAGCAACAAAG | 0.13 | 3MMs | SL3.0ch07:-32490648 | | Intergenic |
| TGGAATTAGAAAAGGCGACAGAG | 0.113 | 3MMs | SL3.0ch01:-31110938 | | Intergenic |
| TGCGCTTAGAGAAAGCGACATGG | 0.112 | 4MMs | SL3.0ch09:+28927574 | gene:Solyc09g031640.1 | CDS |
| TGGAATTGGAGAGAGCCACATGG | 0.105 | 4MMs | SL3.0ch12:+19685978 | gene:Solyc12g077540.2 | CDS |
| TGGAGTTCGAAGAAGTGACAAGG | 0.095 | 4MMs | SL3.0ch11:+38553805 | | Intergenic |
| TGAAATTGGAGGAAGAGAGAGGG | 0.069 | 4MMs | SL3.0ch06:-31118023 | gene:Solyc06g048430.3 | intron |
| TCGATATAGAGGAAGTGACAAGG | 0.067 | 4MMs | SL3.0ch11:-31892066 | | Intergenic |
| TGGAAGTAGAAGAAGCAATAGAG | 0.064 | 4MMs | SL3.0ch04:-22943006 | | Intergenic |
| TGGAATTAGGGGTAGTGACAAGG | 0.051 | 3MMs | SL3.0ch02:+35078725 | gene:Solyc02g062750.3 | intron |

**Supplemental Table S3. qPCR primers**

Table of primers utilized for qPCR to demonstrate silencing in *Nicotiana benthamiana.*

| Primer | Sequence | Details |
|---|---|---|
| Ef1α forward | AGCTTTACCTCCCAAGTCATC | qPCR primer |
| Ef1α reverse | AGAACGCCTGTCAATCTTGG | qPCR primer |
| NbPire1-1 forward | AACGGTTCGATCCCTATTGTG | qPCR primer |
| NbPire1-1 reverse | GTTCCTCCTTCAAGCCTTTG | qPCR primer |
| NbPire1-2 forward | CAGCGGTTCGATCCCTATTG | qPCR primer |
| NbPire1-2 reverse | TTTGGTTCCTCCTTCAAGCC | qPCR primer |
| NbPire1-3 forward | CAATCCCTAACGCTTCCATCC | qPCR primer |
| NbPire1-3 reverse | GCTTGCACTGCTTCAACTTGG | qPCR primer |
| NbPire2-1 forward | ATCGCCAATGCTTCAATCCC | qPCR primer |
| NbPire2-1 reverse | CATTCAATTCCTCATTGGAGCC | qPCR primer |
| NbPire2-2 forward | CCATGCTTTCCTCTAACCCC | qPCR primer |
| NbPire2-2 reverse | TACCCCACCATTCAATTCCTC | qPCR primer |

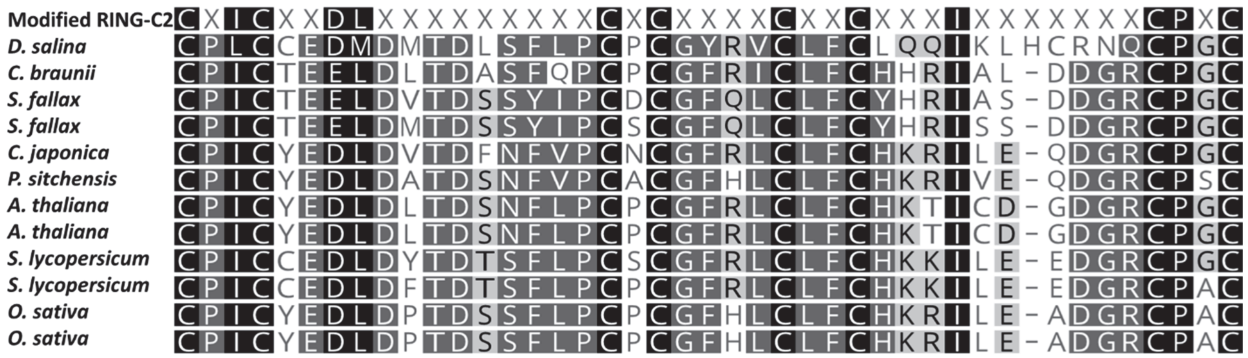

**Supplemental Figure S1. Alignment of the modified RING-C2 domain found in Green Algae, Bryophytes, Gymnosperms, and Angiosperms.**

**(A)** Alignment of a subset of identified modified RING-C2 ubiquitin ligases. The alignment was generated utilizing clustal omega. The top sequence (Modified RING-C2) represents the previously described amino acids found in RING-C2s in Arabidopsis which are important for the interaction with zinc.

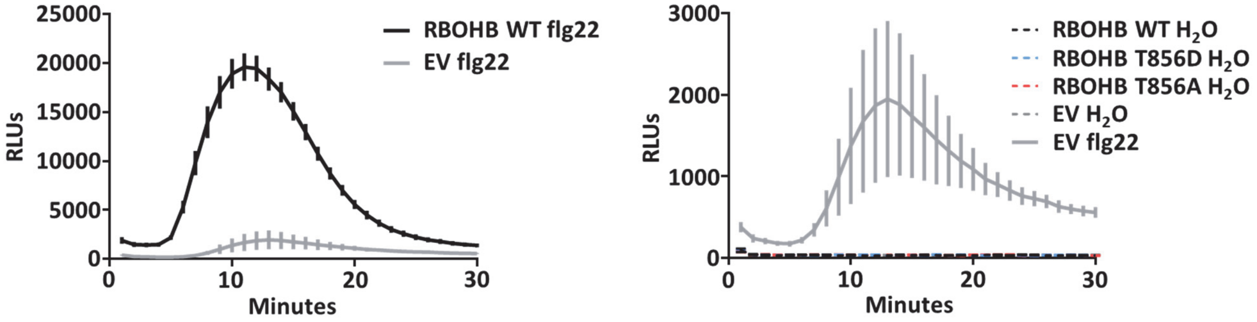

**Supplemental Figure S2. Transient expression of *SlRBOHB* can be differentiated from the endogenous *NbRBOHB* burst after flg22 induction.**

Left Panel: ROS burst is induced in *Nicotiana benthamiana* during flg22 treatment after infiltration with *Agrobacterium* carrying empty vector (EV, solid grey) and the burst is higher after transient expression of RBOHB from *S. lycopersicum* (solid black). Leaf disks were collected from *N. benthamiana* and treated with 100nM flg22 to induce ROS production over 30 minutes. Results display the mean ±SE, n=7 leaf disks, RLU = relative light units. Water controls display no ROS burst. Right Panel: ROS burst after flg22 treatment after infiltration with *Agrobacterium* carrying EV (solid grey). All water treatments did not induce a ROS burst (dotted lines). Note the difference in scale for RLUs between both panels.

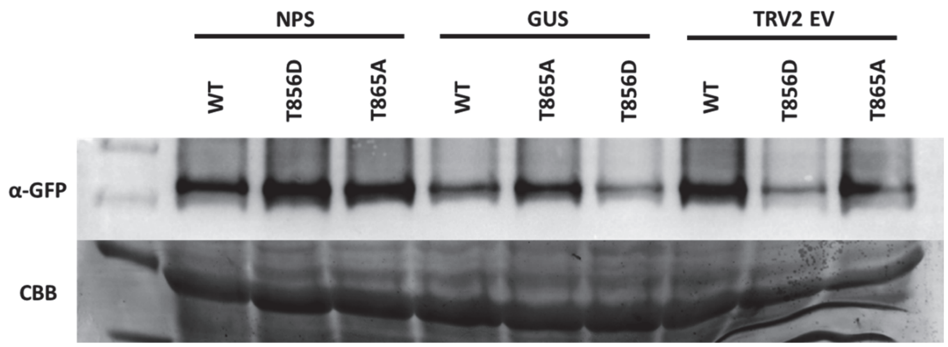

**Supplemental Figure S3. SlRBOHB phosphomutant expression is similar for the GUS and TRV2 controls in *Nicotiana benthamiana.***

Expression levels of SlRBOHB phosphomutants 48h after transient expression in *Nicotiana benthamiana* subjected to virus induced gene silencing (VIGS). Plants silenced for four *N. benthamiana PIRE* homologs (NPS) displayed similar levels of SlRBOHB accumulation. Both GUS and TRV2 empty vector (EV) controls displayed decreased accumulation of $SlRBOHB^{T856}$.

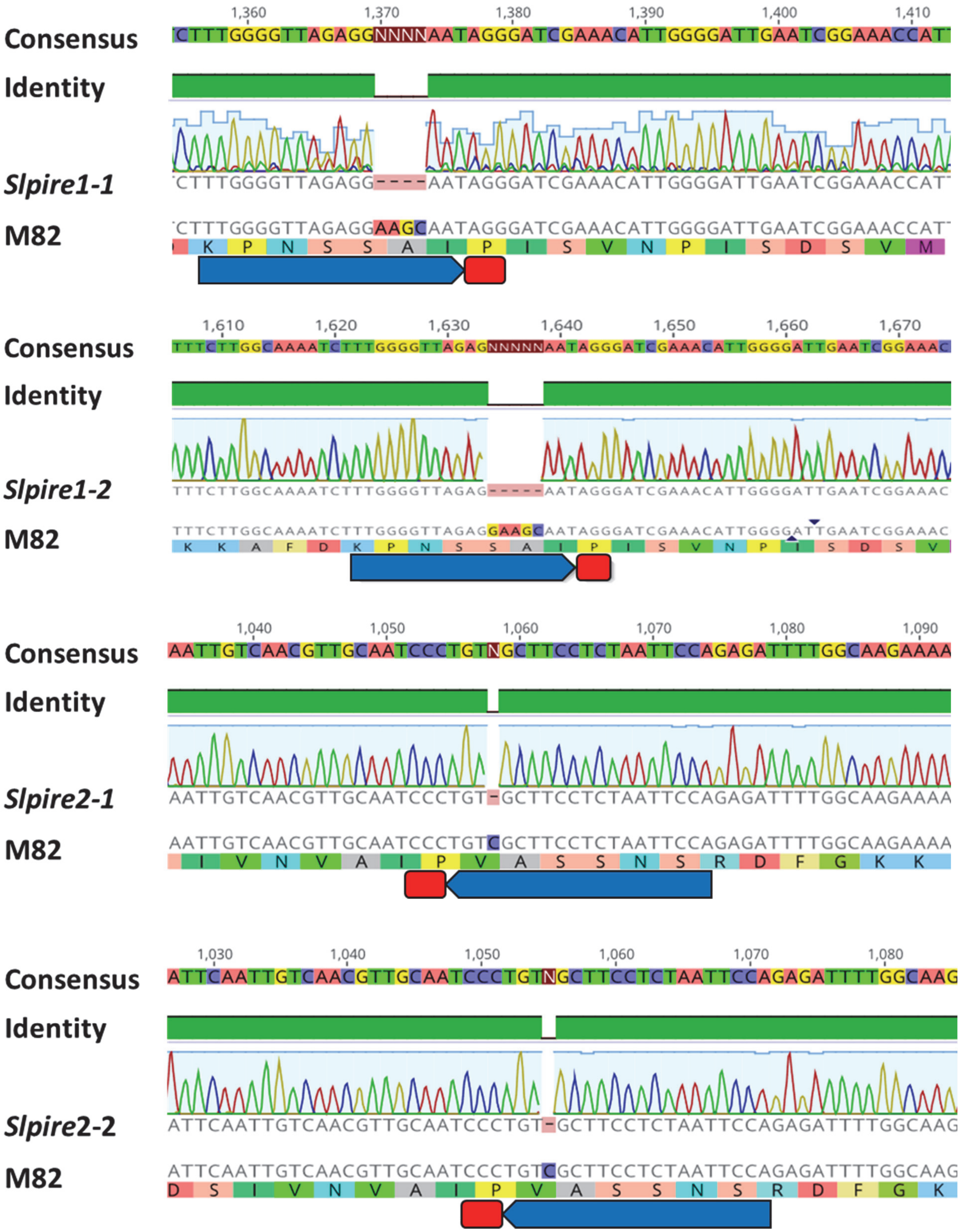

1,360
1,370
1,380
1,390
1,400
1,410
Consensus
CTTTGGGGTTAGAGGNNNNAATAGGGATCGAAACATTGGGGATTGAATCGGAAACCAT
Identity
Slpire1-1
CTTTGGGGTTAGAGG----AATAGGGATCGAAACATTGGGGATTGAATCGGAAACCAT
M82
CTTTGGGGTTAGAGGAAGCAATAGGGATCGAAACATTGGGGATTGAATCGGAAACCAT
K P N S S A I P I S V N P I S D S V M
1,610
1,620
1,630
1,640
1,650
1,660
1,670
Consensus
TTTCTTGGCAAAATCTTTGGGGTTAGAGNNNNNAATAGGGATCGAAACATTGGGGATTGAATCGGAAAC
Identity
Slpire1-2
TTTCTTGGCAAAATCTTTGGGGTTAGAG-----AATAGGGATCGAAACATTGGGGATTGAATCGGAAAC
M82
TTTCTTGGCAAAATCTTTGGGGTTAGAGGAAGCAATAGGGATCGAAACATTGGGGATTGAATCGGAAAC
K K A F D K P N S S A I P I S V N P I S D S V
1,040
1,050
1,060
1,070
1,080
1,090
Consensus
AATTGTCAACGTTGCAATCCCTGTNGCTTCCTCTAATTCCAGAGATTTTGGCAAGAAAA
Identity
Slpire2-1
AATTGTCAACGTTGCAATCCCTGT-GCTTCCTCTAATTCCAGAGATTTTGGCAAGAAAA
M82
AATTGTCAACGTTGCAATCCCTGTCGCTTCCTCTAATTCCAGAGATTTTGGCAAGAAAA
I V N V A I P V A S S N S R D F G K K
1,030
1,040
1,050
1,060
1,070
1,080
Consensus
ATTCAATTGTCAACGTTGCAATCCCTGTNGCTTCCTCTAATTCCAGAGATTTTGGCAAG
Identity
Slpire2-2
ATTCAATTGTCAACGTTGCAATCCCTGT-GCTTCCTCTAATTCCAGAGATTTTGGCAAG
M82
ATTCAATTGTCAACGTTGCAATCCCTGTCGCTTCCTCTAATTCCAGAGATTTTGGCAAG
D S I V N V A I P V A S S N S R D F G K

**Supplemental Figure S4. Validation of *Slpire* gene edited lines by DNA sequencing.** Sequencing results for *Slpire1* and *Slpire2* gene edited lines. Gene editing occurred in the *Solanum lycopersicum* cultivar 'M82'. Blue arrow highlights the gRNA utilized while the red box represents the protospacer adjacent motif (PAM).

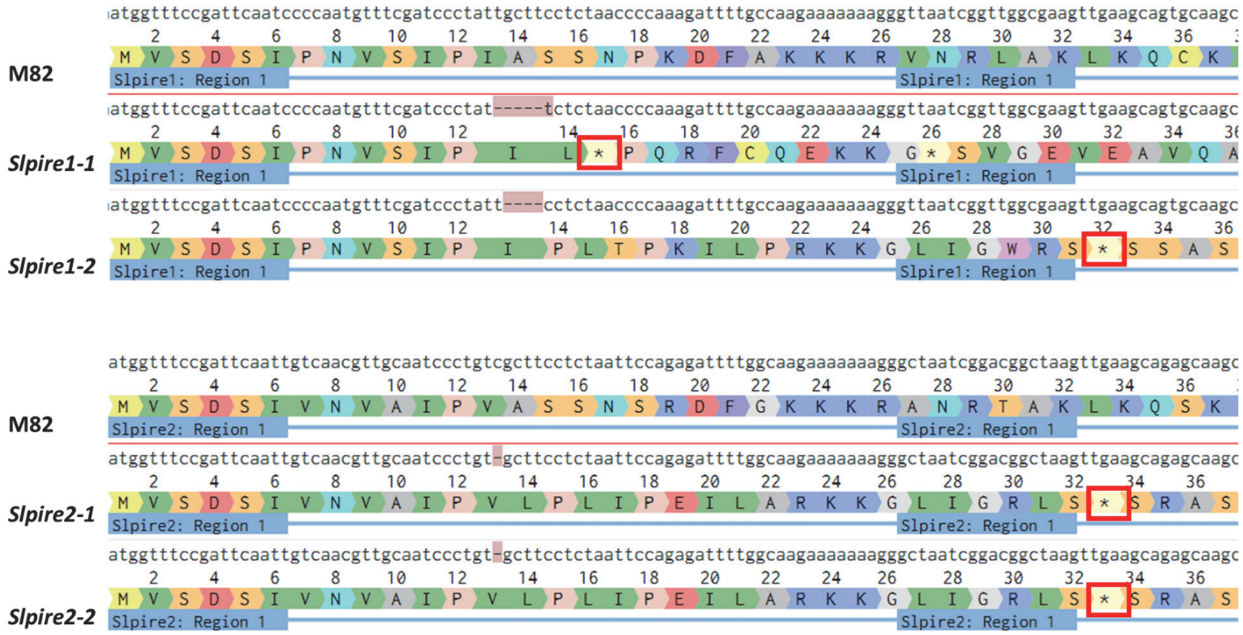

**Supplemental Figure S5. Alignments of the amino acid translations for *Slpire* mutant lines.**

Amino acid alignments displaying the frameshift mutations generating early stop codons on gene edited lines (Stop codons are highlighted by red boxes). Alignment was generated using Clustal omega. Top: *Slpire1* edited lines, bottom: *Slpire2* edited lines.

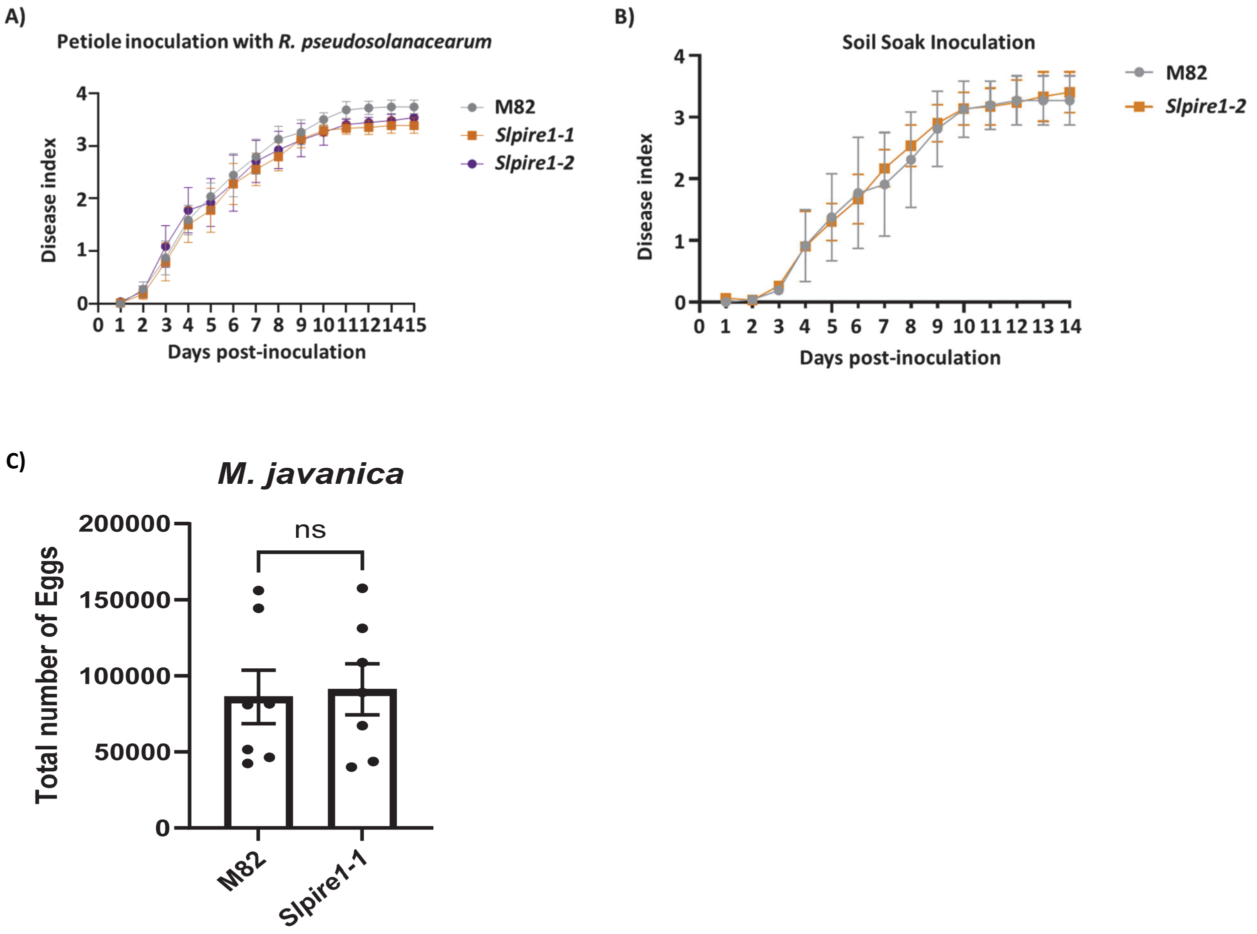

**Supplemental Figure S6. Disease measurements for *R. pseudosolanacearum* GMI1000 strains and *M. javanica* strains on M82 (wild type) and *Slpire1*. A)** The x-axis represents the days post-inoculation, and the y-axis represents the disease index. Error bars correspond to the standard deviation of the Mean (SEM). This experiment was repeated in three biological replicates, and in each replicate N=18 plants were inoculated at the petiole level. **B)** Representative graph of egg counts on infected roots. 4-week-old plants were infected with 500 J2 stage nematodes. Nematode egg counts were performed 7 weeks post infection. Error bars correspond to the standard deviation of the Mean (SEM). This experiment was repeated 2 times with 7 biological replicates per experiment

N=7 plants were inoculated. Statistical differences were detected by t-test, alpha = 0.05, ns = not significant.
